# Differing effects of alcohol use on epigenetic and brain age in adult children of alcoholic parents

**DOI:** 10.1101/2023.09.05.556404

**Authors:** Jamie L. Scholl, Kami Pearson, Kelene A. Fercho, Austin J. Van Asselt, Noah A. Kallsen, Erik. A. Ehli, Kari N. Potter, Kathleen A. Brown-Rice, Gina L. Forster, Lee A. Baugh

## Abstract

It is known that being the adult child of an alcoholic (ACoA) can confer a wide variety of increased health and psychological risks, including higher rates of anxiety, depression, and posttraumatic stress disorder symptoms. Additionally, ACoAs are at greater risk of developing substance use disorders than individuals from non-alcoholic families. To better understand the psychobiological factors underlying these risks, ACoA individuals with risky hazardous alcohol use (n=14) and those not engaged in hazardous use (n=14) were compared to a group of healthy controls. We examined structural brain differences and applied machine learning algorithms to predict biological brain and DNA methylation ages to investigate differences between these groups. Contrary to our hypothesis, we found that hazardous and non-hazardous ACoA groups had lower predicted brain ages than the healthy control group (n=100), which may result from neuro-developmental differences between ACoA groups and controls. When examining specific brain regions, we observed decreased cortical volume within bilateral pars orbitalis and frontal poles, as well as the left middle temporal gyrus and entorhinal cortex within the hazardous alcohol ACoA group, all areas consistent with previous research examining how alcohol use affects brain structure. When looking at the epigenetic aging data, the hazardous ACoA participants had increased predicted epigenetic age difference scores compared to the control group (n=34) and the non-hazardous ACoA participant groups. In summary, the results demonstrate a decreased brain age in the ACoAs compared to control, concurrent with increased epigenetic age specifically in the hazardous ACoA group, laying the foundation for future research to identify individuals that may have an increased susceptibility to developing hazardous alcohol use. Together, these results provide a better understanding of the associations between epigenetic factors, brain structure, and alcohol use disorders.

## 1 Introduction

Hazardous alcohol use, defined as continued alcohol consumption with little or no regard for negative physical or social impacts on one’s life, is an ongoing problem within the United States. The Center for Disease Control and Prevention (CDC) estimates that hazardous alcohol use is a leading cause of preventable deaths in the United States, costing the country over $240 billion annually in combined healthcare costs and loss of productivity (CDC, 2019). Twelve percent of Americans report being raised in a household with at least one parent likely to have an alcohol use disorder (AUD) and meet the criteria for being an adult child of an alcoholic (ACoA; (Grant, 2000)), and that can be as high as 30% among college students (Kelley et al., 2010).

Being an ACoA is associated with many negative outcomes, including higher rates of anxiety, depression, and post-traumatic stress disorder (PTSD) symptoms (Kelley et al., 2010, Klostermann et al., 2011, Woodford et al., 2011), college attrition (Kitsantas et al., 2008), as well as the increased likelihood of the development of substance use disorders compared to individuals from non-alcoholic families (Brown-Rice et al., 2018, Eddie et al., 2015, Yoon et al., 2013). These increased risks are likely due to a combination of genetic and environmental factors (Crane, 2019, Goate and Edenberg, 1998, Köhnke, 2008). Genetic influence can play a role in the response to trauma exposure and in the development of anxiety and depression disorders, possibly due to polymorphisms in related genes. Previous work has shown differences in the 5-HTTLPR alleles (short and long variants) on the SERT gene in hazardous ACoA compared to resilient (Brown-Rice et al., 2018), which has been associated with both depression risk and alcohol dependence (Canli and Lesch, 2007). Relationships involving early life stressors and the short-form of the allele have been reported with a variety of mental health outcomes, including AUD, anxiety, and depression (Kaufman et al., 2007, Stein et al., 2008, Stein et al., 2009, McHugh et al., 2010, Petersen et al., 2012). Individuals that carry the short variant are reported to show deficits in emotional processing and functional differences in the amygdala, insular cortex, and prefrontal cortex, as well as increased activation in the amygdala in response to negative stimuli (Canli and Lesch, 2007, Hariri et al., 2005, Friedel et al., 2009, Stein et al., 2007, von dem Hagen et al., 2011). Additionally, variability in SNPs in cholinergic receptor genes has been related to early alcohol use and increased response to alcohol (Schlaepfer et al., 2008, Joslyn et al., 2008), and genotypes in specific receptor subunits were significantly linked to alcohol use in ACoAs (Brown-Rice et al., 2018). More recently, an assessment of polygenic risk scores has shown associations between AUD, family history of alcohol use, and AUD severity (Lai et al., 2022, Nurnberger Jr et al., 2022).

Several structures in the brain have been associated with changes in both ACoAs (Brown-Rice et al., 2018) and individuals engaged in hazardous alcohol use and AUD (Yang et al., 2016). Structural and functional differences have been found in several brain regions in individuals with a family history of AUD (Cservenka, 2016); specifically, studies have shown greater hippocampal volume in those individuals compared with individuals from non-alcoholic families (Hanson et al., 2010), whereas adolescents and young adults with AUD themselves have been shown to have reduced hippocampal volume compared to controls (Welch et al., 2013). Additionally, previous research has shown increased ventral striatal volume in binge drinkers (common amongst college students) compared to healthy controls but decreased volume in individuals with AUD (Howell et al., 2013). Other reported changes include reduced prefrontal gray matter volume, (Luciana et al., 2013, Yang et al., 2016, Meda et al., 2017) and greater than expected decreases in cortical thickness in adolescents and young adults using alcohol (Luciana et al., 2013, Pennington et al., 2015, Morris et al., 2019), indicating alterations in natural developmental changes that occur in adolescence and young adulthood in those who consume alcohol. Finally, significantly smaller amygdala volume has been seen in alcohol-naïve individuals with a parent who developed AUD by age twenty-five with at least two more first-degree relatives with AUD (Benegal et al., 2007).

However, it is unclear whether these negative outcomes are due to genetic variation, early life stressors experienced by being raised by a caregiver with an alcohol problem, the individual’s own current hazardous alcohol use, or a combination of multiple factors. It is noteworthy that neural functioning can be affected by family history in both resilient and hazardous alcohol use populations (Heitzeg et al., 2008, Heitzeg et al., 2010). Further, not all ACoAs go on to have alcohol problems themselves, which suggests a resilient subpopulation. To better understand the psychobiological factors that underlie hazardous alcohol use in ACoAs, it is necessary to compare ACoA groups with risky or hazardous alcohol use (vulnerable) and groups not engaged in hazardous alcohol use (resilient) against a control group (not ACoA) considered to be representative of the general population.

To examine these relationships, we analyzed structural brain differences between these groups; ACoAs who engage in hazardous alcohol use and those who do not, and an age-matched control group. In recent years, machine learning algorithms have been applied to structural magnetic resonance imaging (MRI) scans to predict a biological brain age (Cole and Franke, 2017). These predicted scores can then be compared to a participant’s chronological age to calculate a difference score or predicted brain age difference (PBAD). Clinically, this PBAD measure has been used in a variety of conditions, including depression (Han et al., 2020), bipolar disorder and schizophrenia (Hajek et al., 2019), and post-traumatic stress disorder (Liang et al., 2019). Typically, psychiatric patients exhibit a higher PBAD than control participants, indicating that these conditions may be associated with accelerated brain aging processes. For the present study, we hypothesized that vulnerable ACoAs would have a higher PBAD than control participants and resilient ACoAs, as has been observed in other psychiatric populations. Further, we predicted these effects would be associated with reduced hippocampal, amygdala, and prefrontal (rostral middle frontal, superior frontal, and inferior frontal regions) volume when compared to resilient ACoAs and healthy controls based on the brain differences previously reported within AUD populations.

Like brain-age prediction techniques, recently, it has been possible to predict one’s chronological age based on patterns of DNA methylation (DNAm). By measuring age-associated methylation changes at specific 5’C-phosphate-G-3’ (CpG) sites, it is possible to create “epigenetic clocks.” These clocks are highly correlated with an individual’s chronological age, with correlation values up to 0.96 (Horvath, 2013). As DNAm is a common epigenetic modification influenced by both genetic and environmental factors, it is also possible to compare epigenetic clock estimates to chronological age, with those individuals with greater epigenetic clock estimates than chronological experiencing age acceleration. Similar to brain age calculations, the predicted EpiAge score can be compared to chronological age to calculate a difference score, predicted EpiAge difference (PEAD). Such measures of age acceleration have been examined with respect to alcohol use in large, demonstrating increased epigenetic age being associated with increased levels of alcohol consumption (Sullivan and Pfefferbaum, 2019, Mavromatis et al., 2022, Martínez-Maldonado et al., 2022, Oblak et al., 2021). Based on these previous results, we predicted that only vulnerable ACoAs would show increased EpiAge when compared to both a control sample and resilient ACoAs.

## 2 Materials and Methods

### 2.1 Participants

All experimental procedures received approval from the Institutional Review Board of the University of South Dakota and were conducted in accordance with the World Medical Association’s Declaration of Helsinki. Twenty-nine participants were recruited and enrolled through advertising at the University of South Dakota as part of a larger study (Brown-Rice et al., 2018). An initial screening was performed to determine eligibility, which included a score of 6 or more out of 30 on the Children of Alcoholics Screening Test (CAST; (Jones, 1983)), which meets the criteria of being an adult child with at least one caregiver that likely has an alcohol use disorder. Written informed consent was obtained from participants, and participants were compensated for their time. Assessment of alcohol use was determined by the Alcohol Use Disorder Identification Test (AUDIT; (Babor, 1992)), where a score of 8 or higher was considered hazardous alcohol use. Twenty-eight participants met all criteria for enrollment, with 14 in each hazardous (mean age = 21 years, range = 18-25 years; 95% Caucasian) and non-hazardous alcohol use (mean age = 21 years, range = 19-24 years; 82% Caucasian) groups. Both groups had a higher percentage of females than males (64% in the hazardous group and 58% in the non-hazardous group). AUDIT scores for the enrolled participants were 15±1.87 for hazardous and 2.8±0.63 for non-hazardous groups; for more detailed explanations of demographics, see Supplementary Table S1 and (Brown-Rice et al., 2018).

#### 2.1.1 Control Participants

In addition to the twenty-eight ACoA participants, a control group was created from data received from one-hundred participants enrolled in previous, unrelated MRI studies conducted at the same university as a representation of the general population. Similar to the ACoA population, this group was made up of 60% female participants with a mean age of 21 years and a range of 18-28 years. A second control group was created from a data set consisting of 34 participants from unrelated studies conducted at the same university with similar demographics for which genetic data were available for epigenetic age analysis. This group was 59% female, with a mean age of 24 years and a range of 18-48 years. All control participants provided written informed consent (Supplementary Table S1).

### 2.2 Psychological and Alcohol Use Measures

The complete measures for this study have been reported elsewhere (Brown-Rice et al., 2018), with details presented here in brief. In addition to the CAST and the AUDIT, assessments included The Beck Anxiety Inventory (BAI; (Beck and Steer, 1993)), the Beck Depression Inventory (BDI-II; (Beck et al., 1996)), and the Posttraumatic Stress Disorder Checklist (PCL; (Weathers et al., 2013)) to determine the presence of anxiety, depression, and post-traumatic stress disorders – all conditions known to affect brain age estimations. Overall, hazardous ACoA participants scored significantly lower in mental, social, general, and perceived health. Additionally, those with hazardous alcohol use reported higher levels of depression and anxiety, as well as higher posttraumatic stress symptoms. For a full description, see (Brown-Rice et al., 2018) and Supplementary Table S1.

### 2.3 Neuroimaging Procedures

#### 2.3.1 MRI Procedures and Blood Sampling

Participants were screened for contraindications to MRI and consented to blood sample collection prior to any scanning. Participants were excluded if they exhibited any contraindications to MRI, possible psychotic or other psychological symptoms, or contraindications to a blood draw such as blood thinners or mastectomy that would make inclusion in the study potentially hazardous to them, resulting in 26 participants completing the blood sampling portion of the study (13 non-hazardous and 13 hazardous). A 5ml blood sample was collected both prior to and immediately after MRI scanning; blood samples were collected in heparin-coated tubes and stored on ice until processing.

All MR Imaging data were collected on a Siemens 3 T Skyra scanner. The acquisition consisted of a 3D anatomical T1 weighted scan, where volumes were collected in the sagittal plane using an MP-RAGE sequence [TR, 1900 ms; TE, 2.13 ms; FOV, 256 × 256× 256; in-plane resolution, 0.9 mm× 0.9375 mm× 0.9375 mm voxels; flip angle, 9°]. Participants laid head-first supine in the bore, with their heads stabilized by noise-canceling headphones and foam padding. Functional MRI volumes were also acquired during various cognitive tasks, the detailed descriptions and findings of which have been reported elsewhere (Brown-Rice et al., 2018).

#### 2.3.2 Structural MRI Analysis

Regional brain volume and cortical thickness were calculated using FreeSurfer software (Version 7.0, http://surfer.nmr.mgh.harvard.edu), a semiautomatic program that analyzes cortical and subcortical regions. The processing pipeline included T1 image motion correction, intensity normalization, transformation to the Talairach space, skull-stripping, segmentation of subcortical structures and gray and white matter, surface tessellation and inflation, and cortical parcellation to map atlases to the spherical registration. The mapping was fine-tuned according to individual anatomy. For the present study, cortical thickness was measured by the shortest distance of the gray/white matter boundary and the cortical surface, along with hippocampal, amygdala, and ventral striatal volume for each individual brain. Quality control steps included visual inspection of all FreeSurfer parcellations, with manual corrections made whenever necessary to ensure correct tissue categorization. The QC pipelines and guidelines developed by the ENIGMA consortium Cortical QC Version 2.0 (http://www.enigma.ini.usc.edu/) were used. Examiners were blind to experimental group at the time of QC implementation. Manual corrections were required in a small (N=4) number of samples, primarily within the control group (N=3), to correctly categorize grey and white matter. No other manual corrections were required or applied. FreeSurfer cortical thickness measurement has been validated against manual measurements (Cardinale et al., 2014).

#### 2.3.3 Voxel-Based Brain Age

Brain age was estimated using the voxel-based brainageR model, which applies a Gaussian process regression model to T1-weighted scans, parcellating gray and white matter to predict chronological age (Cole et al., 2018a). Chronological brain age was subtracted from predicted brain age to create the predicted brain age difference score (PBAD). Therefore, a negative PBAD score indicates a brain age that is estimated to be younger than the participant’s chronological age.

### 2.4 Analysis of DNA Methylation Levels

Blood samples were obtained by standard procedures from non-fasting participants. DNA methylation was assessed using the Illumina Infinium Methylation EPIC Beadchip Array following the manufacturer’s instructions. This platform interrogates over 850,000 methylation sites across the genome at single-nucleotide resolution. Briefly, the isolated DNA is treated with sodium bisulfite, converting unmethylated cytosines into uracil via deamination. Subsequently, methylated cytosines remain unaffected during treatment and are protected from conversion to uracil. Bisulfite conversion was carried out by utilizing the Zymo EZ-96 DNA MethylationTM Kit. Following bisulfite conversion, whole-genome amplification, fragmentation, hybridization, staining, and scanning were carried out according to the protocol. Sample performance, including bisulfite conversion and all internal controls, was assessed by The GenomeStudio Methylation (M) module.

#### 2.4.1 Hannum EpiAge

Epigenetic age was calculated using the Hannum clock (Hannum et al., 2013). This method is based on a set of 71 CpG sites that are largely non-overlapping. The Hannum clock was chosen as it has been previously demonstrated to have high sensitivity to both environmental and cellular aging processes, including inflammatory biomarkers (Irvin et al., 2018) similar to those that would be expected in ACoA populations (Scholl et al., 2022), as well as being sensitive to psychiatric conditions such as post-traumatic stress disorder (Morrison et al., 2019), and socioeconomic position (Hughes et al., 2018). Chronological age was subtracted from predicted brain age to create the predicted epigenetic age difference score (PEAD). Like the PBAD, a negative PEAD score indicates a predicted epigenetic age that is younger than the participant’s chronological age.

### 2.5 Statistical Analysis

All statistical analyses were completed using GraphPad Prism version 9.2.0 for Mac (GraphPad Software, San Diego, California, USA). Effect sizes are reported using omega-square (w^2^).

To confirm there were no baseline differences in chronological age, we examined differences in age between the control and ACoA groups using a one-way analysis of variance (ANOVA) with ACoA status as our independent factor and age as our dependent factor.

Predicted Brain Age Difference (predicted brain age – chronological age; PBAD) was calculated for each participant. As PBAD is known to be overestimated in younger participants and underestimated in older participants, we calculated residualized PBAD scores by regressing chronological age onto PBAD values (Le et al., 2018). A two-way ANOVA with ACoA status (non-hazardous vs. hazardous vs. control) and sex (Male vs. Female) as independent variables, and the residualized predicted brain age difference (PBAD) as the dependent variable was conducted; posthoc comparisons were corrected for multiple comparisons using the Tukey method. Sex was initially included as a factor to ensure there were no systematic prediction differences across males and females.

As this study predicted differences in brain age and that PBAD outcomes give no detail for location in structural differences, additional analyses were performed. To further explore the relationship between predicted brain age and brain structure, supplemental analyses were performed within regions where we expected to find structural differences. Specifically, based on previously reported results, we expected differences within the hippocampal regions, bilateral amygdalae, bilateral cingulate cortex, and orbitofrontal areas of the brain (see Introduction). These additional analyses were run as separate, one-way ANOVAs on the cortical thickness in each brain region in both the right and left brain hemispheres. This resulted in a total of 106 distinct anatomical comparisons (53 left hemisphere and 53 right hemisphere). Due to the number of separate analyses that were run, the Hochberg Step-Up Bonferroni procedure (Hochberg, 1988) was utilized to maintain family-wise error rates at the prescribed .05 level.

Predicted Epigenetic Age Difference (predicted epigenetic age – chronological age; PEAD) was calculated for each participant and was compared using a two-way ANOVA with ACoA status and sex as fixed factors. The main effects and interactions were decomposed using Tukey’s correction for multiple comparisons. Sex was again included as a factor to ensure there were no systematic prediction differences across males and females.

## 3 Results

### 3.1 Effect of ACoA Status on Chronological Age

A one-way ANOVA with ACoA status as the independent factor (non-hazardous vs. hazardous vs. control) and chronological age as the dependent variable revealed no significant main effects (p = .83), with the mean and standard deviations for age being 21.5 +/-2.6 years for the control group, 21.1 +/-1.9 years for the hazardous group, and 21.2 +/-2.6 years for the non-hazardous group.

### 3.2 Impact of ACoA Status on Brain Age

A two-way ANOVA with ACoA status (non-hazardous vs. hazardous vs. control) and sex (Male vs. Female) as independent variables and the residualized PBAD as the dependent variable revealed a significant main effect of ACoA status (F(2,122) = 3.31, p = .040, w^2^ = .035), with those in the ACoA hazardous and non-hazardous groups having a lower PBAD than control participants. However, there was no significant effect of sex, nor a significant sex x ACoA status interaction (p’s >.10). Post-hoc testing of the main effect of ACoA status demonstrated a near-significant difference between the control participants and the hazardous ACoA group (Mean Difference = 2.568, q(125) = 2.994, p = .094). There was no significant difference between the control group and the non-hazardous ACoA group (Mean difference = 2.302, q(125) = 2.684, p = .144), nor between the hazardous and non-hazardous groups (Mean difference = -.2660, q(125) = .2342, p = .985) (See Figure 1). As there were no significant effects of sex found in the brain age analysis, this factor was collapsed in the presented analyses of this dataset.

**Figure 1:**
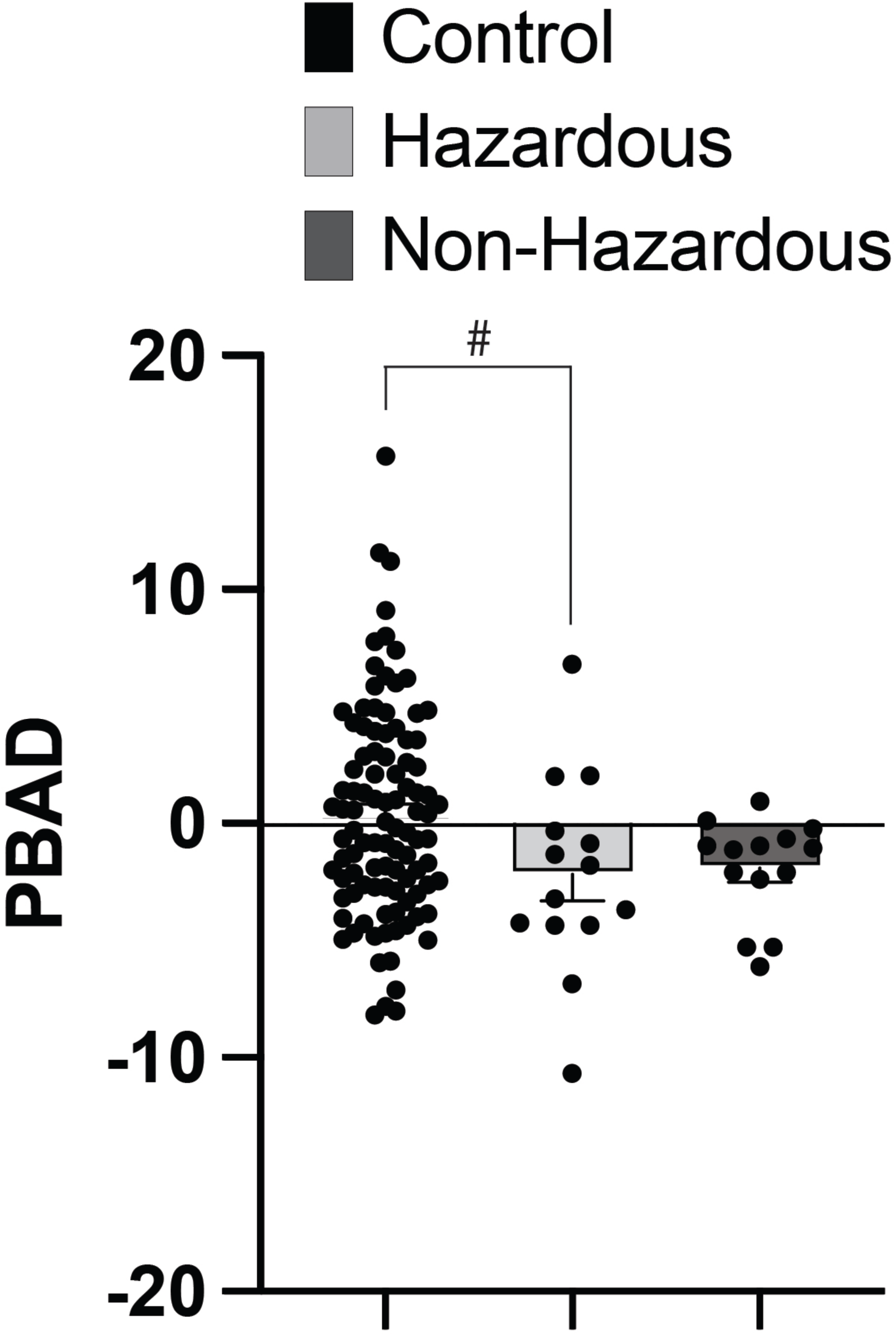
Predicted Brain Age Differences (PBAD) in ACoAs show that, overall, ACoAs have reduced PBAD scores. No significant interaction was observed, with post hoc testing of the main effect of ACoA status showing a marginal difference between the control participants and the hazardous ACoA group. Data represent mean ± SEM. #denotes p < .10.

### 3.3 Impact of ACoA Status on Brain Structure

Next, we explored the effects of ACoA status on each of the previously identified brain regions utilizing separate one-way ANOVAs with each structure’s volume normalized to a per-participant estimated total intracranial volume. This scaling was performed as it is known that certain brain structures scale with general head size (for example, people with larger heads may also have larger hippocampi)(Malone et al., 2015). Analyses were conducted for all 34 regions of the Desikan-Killiany atlas (Desikan et al., 2006) included in each hemisphere’s FreeSurfer automated labeling system. Additionally, as the nuclei of the amygdala and hippocampal subfields (19 anatomical labels per hemisphere) were also of interest, the automated segmentation routine of these structures included in FreeSurfer 7.0 was also used (Iglesias et al., 2015, Saygin et al., 2017, Iglesias et al., 2016). After correcting for the number of comparisons, a total of six regions with significant main effects of ACoA status were observed. These regions with significant values following correction for multiple comparisons consisted of the bilateral frontal pole (Left F(2,125) = 12.69, p < .0001, w^2^ = .155; Right F(2,125) = 6.218, p = .0027, w^2^ = .155), bilateral pars orbitalis (Left F(2,125) = 9.641, p = .0001, w^2^ = .118; Right F(2,125) = 6.766, p = .0016, w^2^ = .083), left middle temporal gyrus (F(2,125) = 4.92, p = .0009, w^2^ = .058), and left entorhinal cortex (F(2,125) = 7.054, p = .0013, w^2^ = .086). Main effects were further explored using Dunnett’s multiple comparisons test to compare each of the ACoA conditions to the control condition (Dunnett, 1955), with significant comparisons illustrated in Figure 2. In all instances where significant differences were observed, they consisted of the %ETIV for the ACoA groups (hazardous and non-hazardous) being significantly smaller than the control group.

**Figure 2:**
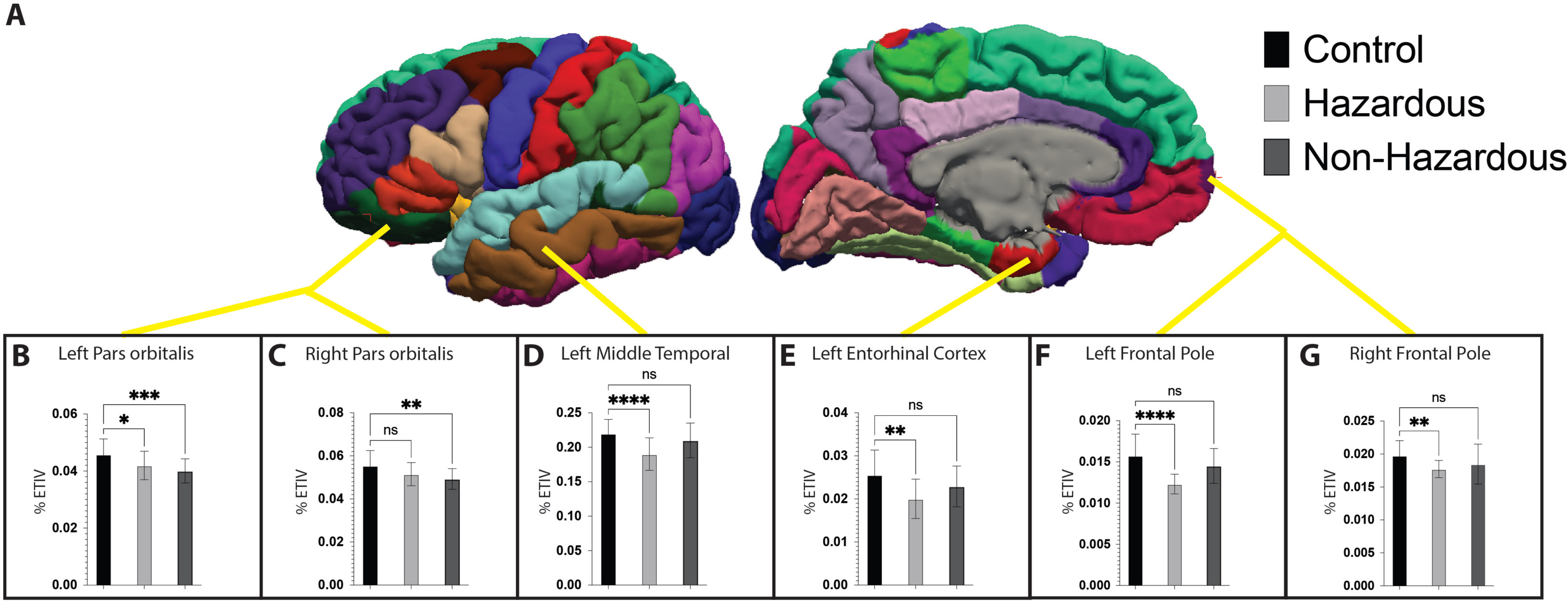
Impact of ACoA Status on Brain Volume. Cortical parcellation utilized the Desikan-Killiany atlas consisting of 34 cortical regions (Figure 2A), as well as nuclei of the amygdala and hippocampal subfields (not shown). Significant main effects of ACoA status were observed for the bilateral frontal pole and pars orbitalis (Figure 2B-D), left entorhinal cortex (2E), and bilateral frontal pole (2F & G). Dunnett’s multiple comparisons were used to test simple effects, with significant comparisons as indicated. Data represent mean ± SEM. *denotes p < .05; **denotes p < .01; ***denotes p < .001; ****denotes p < .0001; ns denotes p > .05.

### 3.4 Impact of ACoA Status on Epigenetic Age

When examining differences in calculated epigenetic age, a significant main effect of ACoA status was observed (F(2,50) = 6.92, p = .002, w^2^ = .183) (See Figure 3). There was no significant effect of sex nor a significant interaction between sex and ACoA status (p’s = .501 and .429, respectively). Post-hoc comparisons revealed a significant difference between the control and the hazardous ACoA group (Mean difference = -3.650, q(50) = 10.74, p = <.0001), as well as between the hazardous and non-hazardous ACoA groups (Mean difference = 2.772, q(50) = 6.478, p = <.0001), characterized by the hazardous condition being associated with an increased PEAD. There was no difference between the control and the non-hazardous ACoA participants (Mean difference = -.8781, q(50) = 2.416, p = .212).

**Figure 3:**
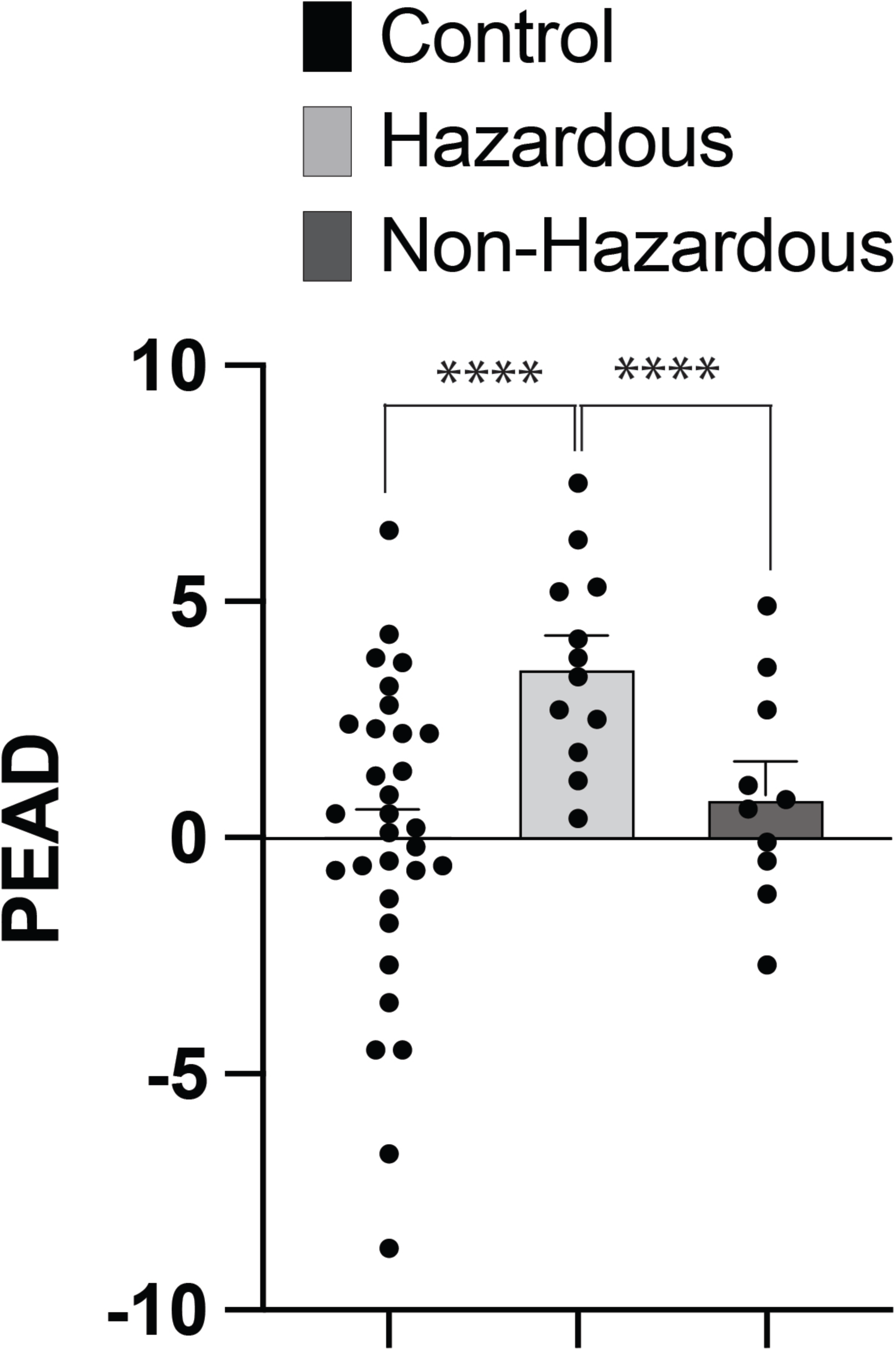
Predicted Epigenetic Age Differences (PEAD) in ACoAs displayed an overall significant main effect of ACoA status, with ACoAs having an increased PEAD score. Post-hoc testing showed a significant difference between the hazardous and control group, as well as the non-hazardous ACoA groups, with the hazardous condition having an increased PEAD score. Data represent mean ± SEM. ****denotes p < .0001.

## 4 Discussion

In this study, we first examined the differences between chronological age and MRI-based predicted brain age in the adult children of alcoholics and compared this difference to a healthy control group. To perform this comparison, we utilized a voxel-based classification method utilizing gaussian process regression, brainageR (Cole et al., 2018a). To better understand the cortical differences reflected in the brain age estimations, we also examined specific cortical regions across each participant group. We chose to examine volume measures, as it is a significant predictor of interest in nearly all brain age models and can be extracted for both cortical and deep-brain structures. Lastly, we examined whether differences between chronological age and epigenetic age varied as a function of ACoA status. To answer this question, we utilized the Hannum epigenetic clock (Hannum et al., 2013) to calculate per-participant epigenetic age estimations.

We predicted that ACoAs that engaged in hazardous alcohol use would have a higher PBAD than both control participants and those ACoAs that were not involved in hazardous alcohol use. Much of this prediction stems from what has been observed both in studies of psychiatric illness (Koutsouleris et al., 2014, Sibille, 2022, Clausen et al., 2022) and what is known about the effects of alcohol on brain structure (see Introduction). We also predicted, based on previous studies in AUD, that much of these differences would be associated with reduced hippocampal, amygdala, and prefrontal volume changes within the ACoA group that engaged in hazardous alcohol use. Lastly, we predicted that, like previous studies examining alcohol use (Sullivan and Pfefferbaum, 2019, Mavromatis et al., 2022, Martínez-Maldonado et al., 2022, Oblak et al., 2021, Luo et al., 2020, Rosen et al., 2018, Fiorito et al., 2019), our ACoA group with hazardous alcohol use would show advanced epigenetic aging reflected in a higher PEAD score than both the control and the ACoA group without hazardous alcohol use.

Testing for the main effect of chronological age on the three ACoA groups revealed no differences. This allowed for the comparison of the effects of ACoA status on predicted age differences, as any observed effects would not be a result of the participant’s chronological age. When examining these predicted age differences (PBAD scores), we found that both hazardous and non-hazardous alcohol use ACoA groups had lower scores than control participants. This result was largely unexpected, as previous studies examining alcohol use disorders have reported an AUD history as resulting in an increased predicted brain age, not a decreased predicted brain age (Ning et al., 2020, Guggenmos et al., 2017). Further, since the participants that were classified as not engaged in hazardous alcohol use ACoAs also demonstrated a decreased brain age, it seems this effect is not entirely driven by current or past alcohol use. When comparisons across ACoA groups were performed, only a significant difference between the control group and the hazardous ACoA group was observed, although the effect approached a significant level between the non-hazardous ACoA group and the control participants (with a p-value of .06).

These differences in the predicted brain age of the ACoA groups may be a result of the age group examined. Importantly, as our study focused on adult children of alcoholics at a mean age of approximately 21 years of age, this is before the maturation of frontal regions is known to have occurred. Although frontal gray matter peaks at approximately 12 years of age (Giedd et al., 1999), it then begins a process of cortical thinning in which frontal cortical gray matter decreases until stabilizing at 20+ years of, at which the ultimate density of frontal cortical gray matter is reached (Giedd et al., 1999, Sowell et al., 2001). It is important to note that this is an age that most of the participants in the present study had not reached, so although all participants were adults, the development of the frontal cortex was likely ongoing. With frontal regions still undergoing maturation, it is possible that the predicted brain age differences observed in the hazardous ACoA group (and approaching significance within the non-hazardous ACoA group) reflect a delayed process of cortical maturation of frontal regions rather than the accelerated brain aging often observed with studies examining alcohol use disorders and psychiatric conditions using brain age estimations in older participant groups. In fact, delayed frontal cortex development specific to regions associated with emotion circuity has recently been demonstrated in children that were abused when compared to those that were not (Keding et al., 2021).

To further examine these observed differences in brain structure, we compared a total of 106 distinct anatomical regions across each of our participant groups. This examination was spurred on both as an attempt to provide additional context to the previously observed brain age differences, as well as guided by previous studies showing that a family history of AUD alone is enough to alter brain function (Brown-Rice et al., 2018), and therefore perhaps structure, of several brain regions. It has previously been demonstrated in a small number of studies that when examining adolescents that can be grouped into high (hazardous) and low (non-hazardous) alcohol use that there are changes in brain structure that can be attributed to both current levels of alcohol use and those that are attributable to a family history of alcohol use, such as the frontal cortex and striatum (Heitzeg et al., 2008, Heitzeg et al., 2010). These results are further substantiated by studies showing that adolescents with a family history of AUD have reduced frontal cortex activity (Cservenka and Nagel, 2012, Mackiewicz Seghete et al., 2013) when completing executive function types of tasks. More recently, our group has shown a complex pattern of increases in some frontal regions (such as the middle frontal gyrus) and decreases in other regions of the brain (such as the posterior cingulate) during functional neuroimaging (Brown-Rice et al., 2018). In the present study, we first observed bilateral decreases in pars orbitalis volume when comparing control participants to ACoAs. Smaller volume and/or cortical thickness measurements within this region have previously been associated with AUD (Pandey et al., 2018) and as a predictor of future problematic alcohol use during adolescence (Squeglia et al., 2017), including those with a family history of alcohol use disorder (Henderson et al., 2018). Interestingly, the developmental trajectory of the pars orbitalis is similar to other frontal regions of the brain and therefore consists of increased cortical thickness during childhood, which then decreases as a product of normal development (Sowell et al., 2003, Sowell et al., 2004), making the observed differences unlikely to be reflective of the decreased brain age reported in the ACoA participant groups, but consistent with previous work examining the effects of alcohol use and familial history of alcohol use in brain development. We also observed a bilateral reduction in the volume of the frontal poles when comparing ACoAs with hazardous alcohol use to healthy controls. This region has also been associated with a decrease in grey matter volume with increased alcohol use (Gröpper et al., 2016, Grodin et al., 2013), making these results consistent with previously reported results in hazardous alcohol use populations. Lastly, we observed decreased grey matter volume in the left middle temporal gyrus and left entorhinal cortex in the hazardous ACoA group when compared to control participants. Decreases in the left entorhinal cortex have been associated with an increased likelihood of relapse during entry into alcohol dependence treatment, along with other components of the mesocorticolimbic reward system that is involved in impulse control, craving, hedonics, and emotional regulation (Cardenas et al., 2011). Therefore, although gray matter reductions are consistent with previous research examining how alcohol use affects brain structure, none of the regions identified in our supplementary analysis provides an explanation as to why PBAD scores are, in fact, lower in these groups. This seemingly conflicting result may be a result of disparate effect sizes that each of the two approaches (ROI vs. Brain Age) requires to determine significant differences between groups. When performing our ROI analysis, the necessity to control for Type-I error rates significantly impacts statistical power, with only the largest effects surviving post hoc corrections. In contrast, as brain age estimations do not perform their calculations as a series of separate statistical tests but rather on the dataset, therefore, the features that result in a reduced predicted brain may not be observable in the ROI analyses. This is especially true with the relatively small sample sizes reported in the present work. Accordingly, this may explain why areas specifically hypothesized to show effects, such as the ventral striatum, amygdala, and hippocampus, did not show significant differences as a function of ACoA status in this study. These hypotheses were based on previous work showing reduced hippocampal volume (Welch et al., 2013), decreased striatum volume in individuals with AUD (Howell et al., 2013), as well as smaller amygdala volume for ACoAs with a parent who developed AUD by age twenty-five with at least two more first-degree relatives with AUD (Benegal et al., 2007). Further work with larger data sets is required to determine how the brain region differences reported here fit into the existing literature.

When examining the epigenetic aging data, we saw that the hazardous ACoA participant group had increased PEAD scores when compared to both the control and non-hazardous ACoA participants. These findings suggest that elevated PEAD scores are related to increased alcohol consumption and not simply being the child of parents with hazardous alcohol use. This result contrasts with what was observed with our brain age predictions, where both ACoA groups showed differences from control participants. This finding is consistent with previous studies showing that increased alcohol consumption is associated with an increased epigenetic age estimation (Luo et al., 2020, Mills et al., 2019, Rosen et al., 2018); however, a family history of alcohol consumption is not. Contrary to previous work showing men have higher epigenetic aging rates than women (Engelbrecht et al., 2022, Kankaanpää et al., 2022, Horvath et al., 2016), we found no significant effect of sex on epigenetic aging. However, this is likely due to the restricted age group used in this study, as all participants were currently attending college.

There are several limitations in the presented study. Firstly, further analyses of the data require increased statistical power. As clinical heterogeneity is always a concern with substance use disorders research, a fact which is further compounded by the large sample size typically required for genetic studies, a more thorough examination which a larger number of participants to confirm the presented results would be beneficial. However, the relatively homogenous sample collected here avoids some of the common issues often reported, such as a broad range of chronological age, varying comorbidities, the presence of psychiatric disorders, and other lifestyle factors that are all likely to influence epigenetic aging and are relatively controlled across our participant groups. One primary advantage of increased sample size would be the opportunity to utilize genome-wide association study methods (GWAS) to examine common variants that may be associated with both the increased epigenetic age and decreased brain age reported, similar to recent research (Luo et al., 2020). However, much larger sample sizes would be required. A second drawback to this study is the cross-sectional nature of the sample collected. As both DNA methylation and brain structure can be affected by both genetic and environmental factors, the presented results are unable to determine whether reported differences within our groups are a result of predisposing factors or consequences of environmental exposure. Future studies may be well-served by adopting a longitudinal approach collecting measures across multiple time points to investigate this relationship between nature and nurture. Additionally, the use of a control population created from data received from previous, unrelated MRI and genetic studies conducted at the same university represents the general population that the ACoA population was recruited from but did not offer the same range of comparisons. Future studies would benefit from a control group undergoing the same assessment and psychological measures as the ACoA group to fully examine the relationships between the familial history of alcohol use and one’s own exposure to alcohol. Lastly, other factors that are known to affect epigenetic aging and brain age estimates should be included as covariates in future studies with larger sample sizes. For instance, physical activity and diet are known to have impacts on both brain age estimation and epigenetic age estimation (Cole et al., 2018b, Bittner et al., 2021, Steffener et al., 2016, Quach et al., 2017, Gensous et al., 2020) and are absent from the presented analysis.

In summary, the presented results indicate decreased brain age in adult children of alcoholics, regardless of hazardous alcohol use that may be a result of disruption in the normal development of the frontal lobe. Additionally, epigenetic aging is only increased in those individuals that report hazardous levels of alcohol intake. These findings warrant further, larger-scale study into the shared mechanisms of decreased brain age in both ACoAs with and without hazardous alcohol use, which may lead to the development of novel methods of identifying those individuals that are susceptible to later developing hazardous alcohol use from those that show similar brain changes but remain resilient.

## 7 Acknowledgments

The authors would like to recognize the assistance of the individuals and volunteers who participated in this research, as well as gratefully acknowledge the work of the radiology staff at Avera Sacred Heart Hospital in Yankton, SD, and the University of South Dakota Human Functional Imaging Core.

## 8 Data Availability Statement

The datasets generated for this study are available upon request to all interested researchers. The raw data supporting the conclusions of this article will be made available by the authors without undue reservation.

