## Supplemental Table 1 for "Differing effects of alcohol use on epigenetic and brain age in adult children of alcoholic parents"

Supplementary Table S1: Assessment measures for adult children of alcoholic parents based on current alcohol status, and control participants from unrelated studies representing a general population.

| Demographic Measure | | Non-Hazardous | Hazardous | Control MRI | Control Genetic |
|---|---|---|---|---|---|
| Gender (%) | Male | 36 | 42 | 40 | 41 |
| | Female | 64 | 58 | 60 | 59 |
| Age (range) | | 19-24 | 18-25 | 18-28 | 18-48 |
| Race (%) | White | 82 | 95 | 88 | 85 |
| | African American | 0 | 0 | 2 | 6 |
| | Native American | 0 | 5 | 1 | 0 |
| | Asian American | 5 | 0 | 6 | 9 |
| | Multiracial | 14 | 0 | 3 | 0 |
| High School Class Size (%) | <50 | 20.00 | 28.57 | *Unknown* | *Unknown* |
| | 50-100 | 6.67 | 28.57 | *Unknown* | *Unknown* |
| | 100-150 | 20.00 | 14.29 | *Unknown* | *Unknown* |
| | >150 | 53.33 | 28.57 | *Unknown* | *Unknown* |
| Diagnosed/Treated Neurological or Psychiatric Disorder (%) | | 20.00 | 21.43 | *0* | *Unknown* |

| Assessment (Range) | Scale | Non-Hazardous | Hazardous | t Value | p Value |
|---|---|---|---|---|---|
| CAST (0-30) | | 19.36±0.99 | 20.00±1.60 | 0.348 | 0.729 |
| AUDIT (0-16+) | | 2.68±0.50 | 15.21±1.51 | -8.337 | *<0.001* |
| Duke Health Profile (0-100) | Physical Health | 73.64±4.03 | 64.74±4.48 | 1.480 | 0.147 |
| | Mental Health | 72.73±5.06 | 52.11±5.05 | 2.868 | *0.007* |
| | Social Health | 71.36±4.43 | 54.74±5.43 | 2.397 | *0.021* |
| | General Health | 72.08±4.10 | 57.48±4.27 | 2.460 | *0.018* |
| | Perceived Health | 77.27±5.43 | 50.00±8.55 | 2.767 | *0.009* |
| | Self Esteem | 68.18±5.45 | 55.26±5.37 | -1.677 | 0.102 |
| | Anxiety | 34.84±5.12 | 49.46±5.13 | -2.006 | 0.052 |
| | Depression | 32.73±5.27 | 48.95±4.64 | 2.275 | *0.029* |
| | Anxiety and Depression | 32.15±4.90 | 51.76±5.44 | -2.686 | *0.011* |
| | Pain | 29.55±6.29 | 47.37±8.09 | 1.762 | 0.086 |
| | Disability | 0.00±0.00 | 15.79±5.48 | -3.108 | *0.004* |
| BDI-II (0-63) | | 11.5±1.94 | 19.53±3.08 | -2.270 | *0.029* |
| BAI (0-63) | | 8.09±1.57 | 13.68±2.15 | -2.136 | *0.039* |
| PCL (17-85) | | 36.64±3.25 | 47.95±3.77 | -2.286 | *0.028* |

*MRI: Magnetic Resonance Imaging, CAST:* Children of Alcoholics Screening Test; AUDIT: Alcohol Use Disorders Identification Test; BDI-II: Beck test for depression; *BAI:* Beck test for anxiety; *PCL:* PTSD Check List.
